# Distinct Alterations in Cerebellar Connectivity with Substantia Nigra and Ventral Tegmental Area in Parkinson’s Disease

**DOI:** 10.1101/2021.05.13.443998

**Authors:** Ian M. O’Shea, Haroon S. Popal, Ingrid R. Olson, Vishnu P. Murty, David V. Smith

**Author notes:** ***Correspondence*** David V. Smith or Vishnu P. Murty Department of Psychology, Temple University. authors contributed equally to this work (Vishnu P. Murty, David V. Smith).

## Abstract

In Parkinson’s disease (PD), neurodegeneration of dopaminergic neurons occurs in the midbrain, specifically targeting the substantia nigra (SN), while leaving the ventral tegmental area (VTA) relatively spared in early phases of the disease. Although the SN and VTA are known to be functionally dissociable in healthy adults, it remains unclear how this dissociation is altered in PD. To examine this issue, we performed a whole-brain analysis to compare functional connectivity in PD to healthy adults using resting-state functional magnetic resonance imaging (rs-fMRI) data compiled from three independent datasets. Our analysis showed that across the sample, the SN had greater connectivity with the precuneus, anterior cingulate gyrus, and areas of the occipital cortex, partially replicating our previous work in healthy young adults. Notably, we also found that, in PD, VTA-right cerebellum connectivity was higher than SN-right cerebellum connectivity, whereas the opposite trend occurred in healthy controls. This double dissociation may reflect a compensatory role of the cerebellum in PD and could provide a potential target for future study and treatment.

## Introduction

The pathological process underlying Parkinson’s Disease (PD) targets dopamine neurons in the midbrain. In early phases of the disease, neurodegeneration occurs in the dopaminergic neurons of the substantia nigra (SN), an area of the midbrain responsible for initiating movement through the nigrostriatal network. However, the dopaminergic neurons in the ventral tegmental area (VTA), a neighboring midbrain structure, are relatively spared from the neuronal degradation in early stages of PD (Dagher & Robbins, 2009; Damier et al., 1999; Fearnley & Lees, 1991, Hirsch, Graybiel, & Agid, 1988). These neurons play a pivotal role in the mesocortical and mesolimbic pathways, functional networks responsible for executive function, reward signaling, and motivation. The lack of degeneration in the VTA may explain why PD patients display motor deficits like bradykinesia and tremors but maintain the ability to engage in motivated behavior (Dagher and Robbins, 2009). This differentiation in behaviors that are affected and remain intact in early PD are thought to result from the discrete patterns of connectivity arising from the SN and VTA areas of the dopaminergic pathways. However, neuroimaging work has yet to investigate the way the SN and VTA differentially interact with the rest of the brain in PD compared to controls.

Within the midbrain, the SN and VTA perform separate yet parallel functions ranging from attention to learning to action (Berridge et al., 2009, Salamone et al., 2007, Wise, 2004). Previous work from our lab provides evidence of reliable differences in two distinct functional networks of the midbrain in controls during resting state (Murty, et al., 2014). The SN had greater connectivity with sensorimotor areas of the cortex like the precentral gyrus while the VTA had greater connectivity to areas associated with reward and motivation such as the nucleus accumbens (NAcc). Notably, other organizational schemas of the midbrain have challenged the notion of treating these two nuclei as distinct, but rather treat them as a unified structure varying across a continuous gradient (Bromberg-Martin et al., 2010; Düzel et al., 2009). However, PD is proposed to affect the SN to a greater extent than the VTA, highlighting a neurological condition that predicts a dissociation between these two networks.

A growing body of work has used resting-state functional magnetic resonance imaging (rs-fMRI) to study midbrain networks in PD, focusing on interactions of the SN with systems responsible for motor planning and execution (Hacker et al., 2012; Wu et al., 2012; Sharman et al., 2013. Previous work found that the SN has decreased functional connectivity to the supplementary motor area, default mode network, and dorsolateral prefrontal cortex in patients with PD, where in controls there is increased SN connectivity to these regions (Wu et al., 2012). Administration of Levodopa partially normalized these differences, indicating dopamine’s role in regulating connectivity (Wu et al., 2012). Others suggest that SN connectivity is reduced to the thalamus, globus pallidus, and the putamen (Sharman et al., 2013). Further investigation of the striatum indicated decreased striatal connectivity with the midbrain, however the SN and VTA were not studied individually (Hacker et al., 2012). This research with rs-fMRI focused on larger networks that contribute to motor functioning without examining the more granular interactions of sub-regions. Namely, this work did not include the VTA and its projections to mesocortical and mesolimbic systems, thus leaving open questions about how interactions between the SN and VTA may be altered by changes in dopaminergic tone.

The goal of the present study was to investigate dissociations between dopaminergic midbrain networks in patients with PD and healthy controls. We predicted that the VTA connectivity with cognitive regions of the NAcc and subgenual cingulate will not be different across groups, whereas, SN connectivity with the putamen, supplementary motor area, and primary motor cortex will be greater in controls than PD patients. The few studies that have examined related questions used relatively small sample sizes (Hacker et al., 2012; Wu et al., 2012; Sharman et al., 2013), which may affect reproducibility of findings. Thus for the present study, we combined three open datasets of rs-fMRI data in order to obtain robust results. We first conducted a whole-brain analysis to analyze functional connectivity to the SN and VTA, which were defined by probabilistic atlases (Murty et al., 2014). Connectivity values were compared across these seed regions in whole-brain analyses to examine the differences between the two networks. We focused on the group by region of interest (ROI) interaction for network-specific effects of PD. To preview our findings, we find that PD differentially affects the two dopaminergic networks in the midbrain.

## Results

First, we wanted to determine the differences in functional connectivity of the SN compared to the VTA, collapsing across group. Whole-brain analysis showed a significant effect of ROI (p < 0.05, whole-brain corrected), such that the SN had greater connectivity than the VTA to various regions throughout the cortex, including the precuneus, anterior cingulate gyrus, and areas of the occipital cortex, partially replicating previous results (Figure 1; Table 1; Murty et al., 2014). However, unlike prior reports in younger adult populations, the reverse contrast of VTA greater than SN did not show any significant differences at our correction threshold.

**Figure 1.**
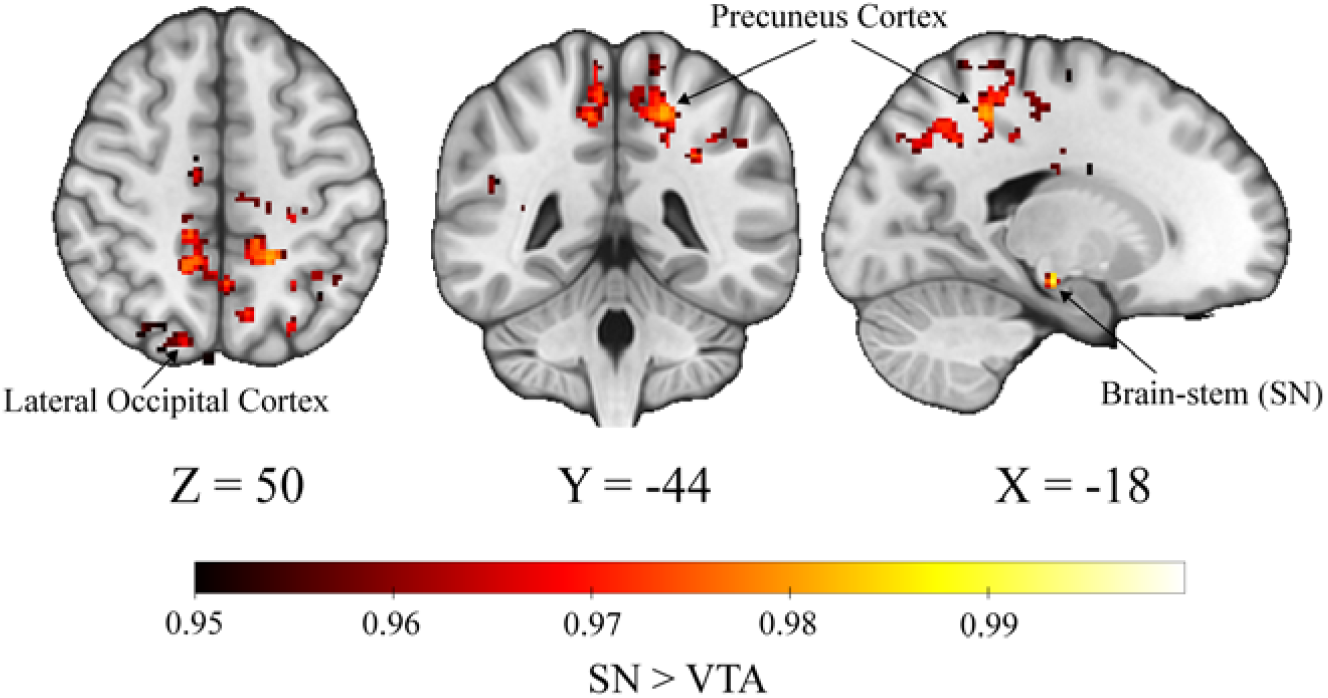
Whole-brain connectivity analysis reveals a main effect of SN>VTA. Image is thresholded at p < 0.05 Coordinates correspond with peak activation of the largest cluster, in the precuneus cortex.

**Table 1.** Voxel clusters where connectivity is greater to SN than VTA. The table lists region name, amount of voxels, peak activation, and coordinates in MNI space for clusters significant at a p < 0.05 threshold. Clusters of less than 10 voxels (8 clusters) were removed from the table.

| Region<br>SN > VTA | Cluster<br>(voxels) | Peak<br>Activation | x | y | z |
| --- | --- | --- | --- | --- | --- |
| Precuneus Cortex (Left) | 2056 | 0.981 | -18 | -44 | 50 |
| Brain-stem (right SN) | 291 | 1 | 12 | -24 | -20 |
| Brain-stem (left SN) | 241 | 1 | -10 | -20 | -16 |
| Cingulate Gyrus, anterior division (left) | 235 | 0.981 | -4 | -8 | 30 |
| Lateral Occipital Cortex, superior division (right) | 173 | 0.973 | 18 | -76 | 54 |
| Supramarginal Gyrus, posterior division/<br>Angular Gyrus (right) | 100 | 0.967 | 48 | -42 | 24 |
| Supramarginal Gyrus, anterior division (right) | 65 | 0.964 | 68 | -26 | 34 |
| Angular Gyrus (left) | 53 | 0.962 | -44 | -56 | 16 |
| Lateral Occipital Cortex, superior division (left) | 34 | 0.955 | -32 | -60 | 54 |
| Cerebral White Matter (left) | 20 | 0.964 | -16 | -14 | 28 |
| Lateral Occipital Cortex, superior division (right) | 12 | 0.954 | 46 | -72 | 18 |
| Superior Frontal Gyrus (left) | 12 | 0.955 | -14 | -10 | 68 |
| Cingulate Gyrus, anterior division (right) | 10 | 0.956 | 6 | 4 | 30 |
| Lateral Occipital Cortex, superior division (left) | 10 | 0.953 | -26 | -64 | 46 |

Next, we wanted to determine differences in connectivity between PD patients and the healthy controls, collapsing across seed ROIs. Again, a whole-brain analysis was performed to identify regions whose connectivity differed across the two groups. There were no main effects of group, such that collapsed across ROI, functional connectivity did not differ when comparing the PD group to the controls.

Finally, we wanted to determine if there were any significant interactions between group and ROI to investigate whether PD had a region-specific effect on functional connectivity. Whole-brain analysis was performed, which indicated a significant group by ROI interaction in the right cerebellum (p<0.05, whole-brain corrected; Figure 2A-B). Post-hoc analysis using a paired t-test comparing VTA and SN connectivity within each group revealed that in PD patients, the VTA had enhanced functional connectivity with a 21-voxel region in the right cerebellar cortex (e.g. VIIB and VIIIA) compared to the SN (t = -5.20, p < 0.001, 95% CI = [-0.80, -0.36]), while in healthy controls there was greater SN than VTA connectivity with the right cerebellar cortex (t = 4.69, p < 0.001, 95% CI = [0.31, 0.77]; Figure 2C). In the SN, contralateral and ipsilateral connectivity to the right cerebellar cortex were not significantly different. Effects of laterality were not investigated in the VTA due to its medial location in the brain. To assess laterality of midbrain connectivity in the cerebellum, we left-right flipped the cerebellar region and compared it directly to our original image. We found that both left and right cerebellar regions exhibit a similar pattern of double dissociations in connectivity. However, we found a three-way interaction between seed ROI, group, and cerebellar hemisphere (left/right), such that the ROI * group effect was stronger in the right cerebellum compared to the left (F(1, 404) = 12.4, p < 0.001; Supplementary figure 1).

**Figure 2.**
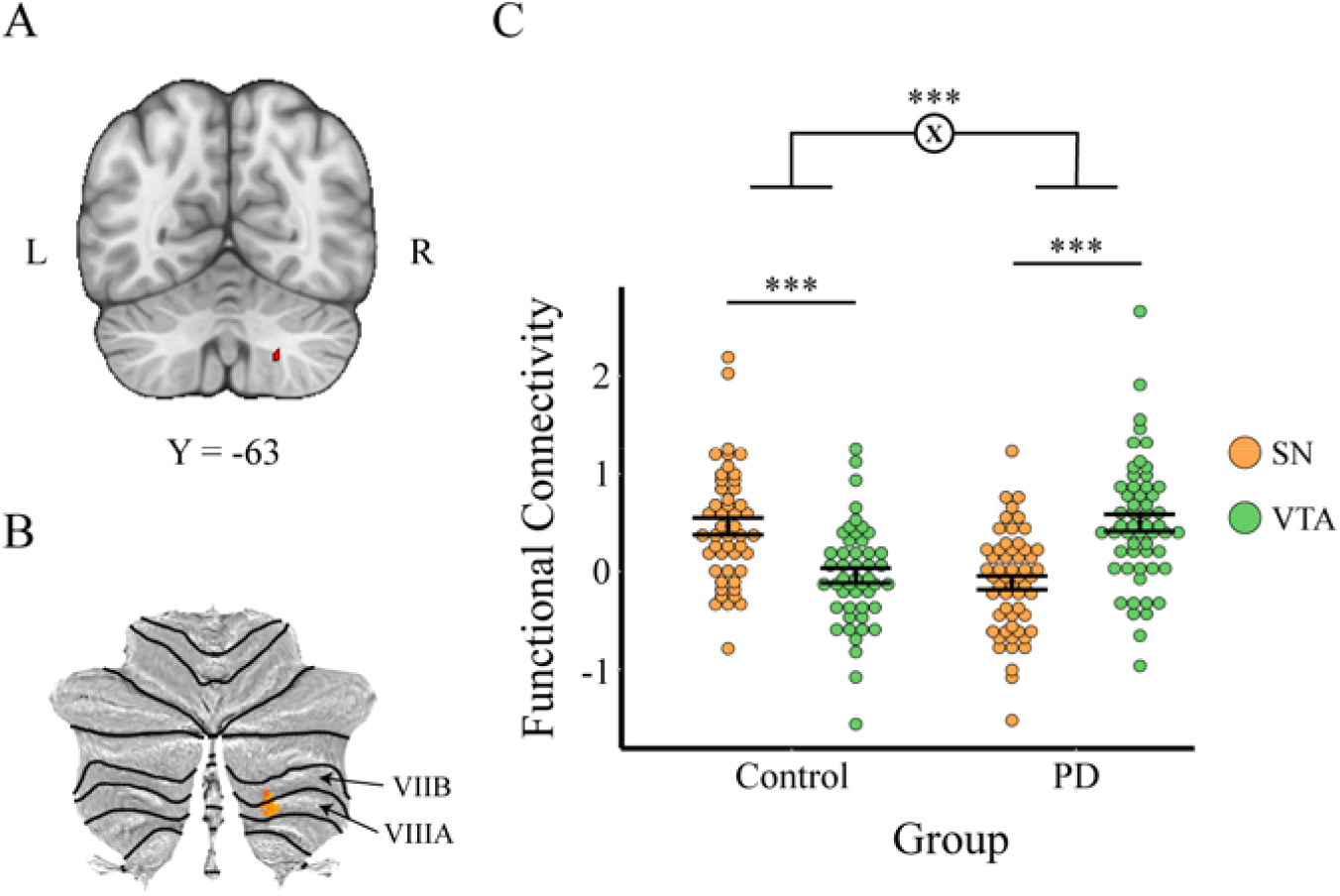
Whole-brain connectivity analysis reveals group by ROI interaction in right cerebellum. A. Axial view of brain showing 21-voxel cerebellar cluster derived using TFCE. Image thresholded at p < 0.05. B. The same cluster as in A., but flat-mapped, shows that the region corresponds to VIIB and VIIIA regions of the cerebellum; and C. Dotplot reveals direction of interaction, where the PD group had greater VTA than SN connectivity with the right cerebellum and the control group had greater SN than VTA connectivity to the right cerebellum. (p < 0.001 ***). Each dot represents one subject’s t-statistic for connectivity, color coded for each specific ROI and divided into control and PD subjects.

Although our primary analyses controlled for differences across datasets using established methods (Friedman & Glover, 2006), we performed additional control analyses to confirm that the observed interaction was not driven by a single dataset. To rule out this confound, we extracted the cerebellar connectivity within each condition and dataset and conducted three 2×2 ANOVAs. In each ANOVA, the interaction was significant (Dataset 1: F(1,72) = 24.2 p < 0.001; Dataset 2: F(1,72) = 22.4, p < 0.001) or approaching significance (Dataset 3: F(1,50) = 3.18, p = 0.08), and the effects were in the same direction. The consistent findings across the three datasets provides confidence in our findings.

Following our original analysis, we were interested in any interaction of the VTA and SN with striatal regions. To investigate any differences in connectivity, we calculated average connectivity of VTA and SN with striatal regions including the caudate, putamen, and NAcc Using a 2 * 2 ANOVA, we found no effects of seed ROI (caudate: F(1, 202) = 0.47, p = 0.493; putamen: F(1, 202) = 1.24, p = 0.266, NAcc: F(1, 202) = 0.78 p = 0.381), group (caudate: F(1, 202) = 0.04 p = 0.842; putamen: F(1, 202) = 0.66 p = 0.418, NAcc: F(1, 202) = 0 p = 0.996), or their interaction (caudate: F(1, 202) = 1.20, p = 0.275; putamen: F(1, 202) = 1.96, p = 0.163, NAcc: F(1, 202) = 2.08, p = 0.151) on connectivity of the midbrain with any striatal regions.

## Discussion

Using three publicly available rs-fMRI datasets, we provide novel evidence of differential connectivity of the SN and VTA in Parkinson’s disease. Namely, we find that SN functional connectivity with motor and pre-motor cortical networks remains intact across PD and control groups. However, we were surprised to find a prominent double disassociation of the midbrain connectivity with the right cerebellum, such that in Parkinson’s, there was greater VTA-right cerebellum connectivity as compared to the SN-right cerebellum—while in healthy controls, there was greater SN-right cerebellum connectivity as compared to VTA-right cerebellum. Together, these findings suggest an important differentiation of connectivity of the dopaminergic midbrain with the cerebellum, when taking into account functional heterogeneity across the midbrain.

Although the cerebellar findings were unexpected, they are not without precedent. Projections from the deep cerebellar nuclei to the VTA are monosynaptic and bidirectional. D’Ambra and colleagues (2021) showed that microstimulation of the deep cerebellar nuclei in mice modulated NAcc spiking activity, both excitatory and inhibitory, depending on the particular location recorded from within the NAcc (D’Ambra et al., 2021). Another study found that optogenetic stimulation of cerebellar-VTA axons dramatically altered goal-oriented behavior (Carta et al., 2019). Older rodent studies indicate the presence of both dopaminergic and non-dopaminergic projections from the VTA to cerebellar cortex, terminating in both Crus I and Crus II (Ikai et al., 1992). In humans, this circuit is considered to be part of a larger, highly integrated learning system (Caligiore et al., 2019). Structural connectivity work in humans shows extensive connections between the basal ganglia and cerebellum, including a pathway between the SN and dentate nucleus of the cerebellum (Milardi et al., 2016). The anatomical connections between structures of the midbrain and cerebellum provide potential pathways by which this differentiation arises, and here we extend this work to show that this is a functional circuit that is altered in PD.

In PD, the role of the cerebellum is relatively unclear, however Wu & Hallett (2013) suggest that the structure has both pathological and compensatory effects. These compensatory effects are thought to help maintain both motor and cognitive function to make up for the degeneration of dopaminergic neurons in the SN (Wu & Hallett, 2013). Therefore, increased VTA-cerebellar connectivity may be the result of the cerebellum overcompensating for neurodegeneration in the SN, while the relatively decreased SN-cerebellum connectivity in PD could be indicative of a pathological effect. In line with this interpretation, reward magnitude has been shown to correlate with cerebellar activity in PD, while in controls, reward magnitude correlates with activity of prefrontal, rhinal cortices and thalamic activity (Goerendt et al., 2004). Thus, these findings correspond with motivated learning being mostly spared in PD while motor function deteriorates (Dagher and Robbins, 2009). It should however be noted that functional connectivity cannot accurately assess the directionality of VTA-cerebellar connections. Overall, we corroborate previous findings that the cerebellum may be mediating cognitive function in PD through its interactions across multiple networks in the brain (Gratton et al., 2019; Palmer et al., 2021; Zhang et al., 2016). Importantly, our interpretation is limited to results in resting-state networks. Future work should investigate these possible compensatory mechanisms with behavioral manipulations that elicit motor and reward responses in the scanner.

The specificity of midbrain-cerebellar connectivity may give insight into the functional implications of our findings. Dissociations in midbrain connectivity with the cerebellum spanned cerebellar lobules VIIB and VIIIA. Based on the functional boundaries defined by King et al. (2019), these clusters correspond to executive and attentionally functional regions of the cerebellum (King et al., 2019). Similarly, using LittleBrain, we found that the clusters correspond to cerebellar voxels that project to dorsal and ventral attention networks at rest, reflecting the role of these cerebellar regions in cognition, not motor processes (Guell et al., 2019). Additional work by Stoodley and colleagues suggests that areas VIIB and VIIIA within the right cerebellum are involved in language production, a cognitive function that is typically impaired in PD and is related to cognitive decline (Stoodley et al., 2012; Altmann & Troche, 2011). This provides further evidence that this cerebellar region could be influencing cognitive functions in PD. This is corroborated by findings that the majority of the cerebellum, including the areas in question, maps onto association areas in the cortex (Buckner, 2013). However, this work also presents contrasting results indicating area VIIIA is involved in sensorimotor functions such as tapping (Stoodley et al., 2012). In order to parse out differences in cerebellar functional topography, future research should be conducted in a clinical sample with PD. Given that executive functioning is spared in the early phases of PD, cerebellar engagement with the VTA may represent how the cerebellum maintains executive functioning through a goal-oriented compensatory mechanism. However, future studies relating these connectivity profiles with executive function deficits are necessary to confirm these hypotheses. Furthermore, late-stage executive deficits in PD may reflect the inability of the cerebellum to keep up with dopaminergic degradation. Ultimately, these results indicate the cerebellum as a possible target of treatment for PD in the future and support further investigation into how cerebellar subregions are impacted by dopaminergic degeneration.

Our findings failed to replicate many of the findings previously shown regarding SN connectivity using rs-fMRI in PD. Namely, we did not find deficits in SN connectivity to primary or supplementary motor areas, dorsolateral prefrontal cortex, thalamus, default mode network nodes, or striatal structures in PD, as have previous groups (Hacker et al., 2012; Sharman et al., 2013; Wu et al., 2012; Wei et al., 2018). In fact, we found a similar pattern of connectivity of the SN when compared to the VTA with motor and pre-motor cortical regions that did not differ across groups. One factor that could explain this lack of replication is spatial resolution. Relative to prior studies of rs-fMRI in PD, our spatial resolution was poor, potentially blurring other signals with the VTA and SN and contributing to a Type II error (i.e., false negative) finding where connectivity differences are missed in the analysis. Using higher resolution data in the future may make the SN and VTA more easily dissociable and provide more specificity in the results. However, it is important to note that our mega-analysis provides more power to assess these relationships and may highlight that these prior reports may potentially represent false positives.

Another factor that could explain this lack of replication is medication status. Wu and colleagues (2012) showed that deficits between the SN and cortical targets in motor and executive regions were ameliorated with Levodopa administration (Wu et al., 2012). In our study, we collapsed across three datasets in which PD patients were un-medicated or medicated with Levodopa. Medication status was not evenly distributed across our samples, and thus we could not effectively control for this factor in our analysis. Despite controlling for dataset as a confound in our GLM, these medication differences may have influenced results; specifically, Levodopa administration may have compensated for any SN connectivity deficits. Similarly, information about patient motor functioning was not available for all subjects and could not be accounted for in our analysis. Future studies should be designed to compare midbrain-cerebellar connectivity in age-matched samples of healthy controls and individuals with PD. This work should also make sure to specifically control for medication status of the subjects, possibly determining the effects of Levodopa on these connectivity differences. Additionally, it may be necessary to use higher resolution imaging data to determine specificity of relevant regions.

Outside of group-related differences, when collapsing across the entire sample, SN connectivity was higher than VTA connectivity to the precuneus, anterior cingulate gyrus, and portions of the occipital cortex. This finding, in part, replicates SN connectivity from a previous study in our lab conducted in a younger adult population (Murty et al., 2014). Unlike our prior study, however, the VTA did not show enhanced connectivity with the NAcc and subgenual cingulate when compared to the SN, conflicting with previous findings. One reason we may have failed to replicate these findings may be the differences in age across samples. Our original study was conducted in a sample of younger adults aged 18 to 25 (*M* = 21.9), whereas our current sample represents older adults aged 36 to 86 (*M* = 65.87, *SD* = 8.98). As an exploratory analysis, we tested correlations between age and midbrain connectivity in our sample. We found a trending effect such that older participants had decreased connectivity between the SN and caudate (r = -0.18, p = 0.066; Supplementary figure 2). We believe this partial replication of previous studies may be explained by age related changes SN functionality. Prior research across rodents and humans has shown decreases in dopaminergic neuromodulation throughout aging, which could alter the connectivity of the VTA at the population level (Bäckman et al., 2006, Dreher et al., 2008). We do replicate findings of strong positive SN connectivity to the supramarginal gyrus and anterior cingulate, however, we also find positive connectivity with the precuneus, occipital cortices and angular gyrus, conflicting with the findings from previous studies (Tomasi & Volkow, 2014). Similar to findings in DTI work we also find greater SN than VTA connectivity as determined by fiber tracking across multiple areas of the cortex, but fail to replicate subcortical findings (Kwon & Jang, 2014). It should also be noted that the SN did not demonstrate greater functional connectivity to regions of the cerebellum as previous results have indicated (Zhang et al., 2015). Given these mixed results, future studies with improved resolution and larger, more diversified samples will be needed to further delineate SN and VTA networks.

Overall, we provide corroborating evidence that there are two dissociable midbrain networks, stemming from the SN and VTA, respectively. Not only does the dissociation provide a better understanding of PD, it also implies that these two regions contribute to cognition and behavior differently via their distinct connections throughout the brain. This observation has implications for other disorders like schizophrenia, substance abuse, ADHD, and depression, disorders in which the pathophysiology is related to dopamine dysfunction. We also provide evidence that the cerebellum may play a significant role in regulating these pathways, consistent with recent work in mice (D’Ambra et al., 2021) and therefore in the development of neuropsychiatric conditions. A growing body of work links mood disorders such as depression to dysregulated functional connectivity within the cerebellum (Frazier et al., 2021; Tepfer et al., 2021). Of note, depression is common in PD (Reijnders et al., 2008); whether this can be linked to common pathways is a question for future research. Overall, our study provides a foundation for future translational work by furthering our understanding of the midbrain’s role in cognition and motor control and characterizing how dysregulated cerebellar connectivity contributes to PD.

## Methods

### Datasets

Resting state functional magnetic resonance imaging (rs-fMRI) data were aggregated from three publicly available sources: 27 PD patients and 16 healthy controls from the NEUROCON project (Badea, et al., 2017), 20 PD patients and 20 controls from the Tao Wu group (Badea, et al., 2017), and 14 de novo PD patients and 14 controls from Tessa and colleagues. (Tessa, et al., 2019). In total, there were 111 participants: 61 PD patients and 50 healthy controls (Table 2). We note that eight subjects were removed from this sample based on image quality metrics as described in the neuroimaging preprocessing section. The final sample consisted of 103 participants: 55 PD patients (Aged 36 – 86, *M* = 65.92, *SD* = 8.90) and 48 healthy controls (Aged 38 – 82, *M* = 65.81, *SD* = 9.12). We note that there are no significant differences in age (t(98) = 0.063, p = 0.95) or sex distribution (chi-square = 0.94, p = 0.33) across PD patients and controls.

**Table 2.** Patient characteristics for each dataset. The Hoehn and Yahr (H&Y) Scale measures the stage of functional disability in Parkinson’s, where 1 is least severe and 5 is most severe. N/A refers to information that was not available from authors of the datasets.

| PD Patient Characteristics | Dataset 1<br>(Badea et al., 2017) | Dataset 2<br>(Badea et al., 2017) | Dataset 3<br>(Tessa et al., 2019) |
| --- | --- | --- | --- |
| Proportion Medicated | 0.926 | N/A | 0 |
| H&Y (SD) | 1.93 (0.33) | 1.88 (0.63) | N/A |
| Age (SD) | 68.7 (10.6) | 65.2 (4.4) | 63.7 (11.1) |

### Pre-registration

The study was pre-registered using AsPredicted (https://aspredicted.org/blind.php?x=vs5zn6). There were slight deviations in our methods from the pre-registration. Using whole-brain analysis deviated from the pre-registration, which indicated that an ROI-based analysis would be used. Given that this sample was made up of older adults—some of whom were diagnosed with PD—there may be different localizations of cortical ROIs due to cortical atrophy or thinning. Therefore the cortical ROIs from previous work in young adults were not used, and instead a whole-brain analysis was performed to ensure that no areas of significant connectivity were missed in the analysis.

### Image Acquisition

For the three datasets, there were various deviations in image acquisition (Table 1). Notably, datasets 1 and 3 were collected using a 1.5 Telsa Siemens Magneto Avanto MRI scanner, while dataset 2 was collected using a 3 Tesla Siemens Magnetom Trio scanner. All anatomical images were collected using a magnetization prepared rapid gradient echo sequence (MPRAGE), and all functional images were collected using echo planar imaging (EPI) sequences. Dataset 3 specifically used T2-weighted EPI sequences with interleaved slice acquisition.

Although there are some notable acquisition differences across these three datasets (Table 3), our analyses explicitly control for these differences (as detailed below). Moreover, such inter-dataset differences should not necessarily lead to false positives, given the distributions of PD patients and healthy controls. Nevertheless, we acknowledge inter-dataset differences could contribute noise to our analyses, potentially increasing the risk of false negatives, but also enhance the generalizability of our results across datasets. We also acknowledge that the relatively coarse spatial resolution could blur signal from adjacent anatomical areas. However, other neuroimaging studies have used relatively similar in-plain axial slice resolution (∼3-4 mm) to characterize these midbrain structures (Adcock et al., 2006; Mallol et al., 2008; Murty et al., 2011; Shohamy & Wagner, 2007; Wu et al., 2012).

**Table 3.** fMRI acquisition parameters for the three datasets. N/A refers to information that was not available from authors of the datasets.

| fMRI Acquisition Parameters | Dataset 1<br>(Badea et al., 2017) | Dataset 2<br>(Badea et al., 2017) | Dataset 3<br>(Tessa et al., 2019) |
| --- | --- | --- | --- |
| **Anatomical** |  |  |  |
| Repetition time (TR) | 1940 ms | 1100 ms | 1900 ms |
| Echo time (TE) | 3.08 ms | 3.39 ms | 3.44 ms |
| Voxel size | 0.97 x 0.97 x 1 mm | 1 x 1 x 1 mm | 0.859 x 0.859 x 0.86 mm |
| Inversion time | N/A | N/A | 1100 ms |
| Field of view (FOV) | N/A | N/A | 220 x 220 mm |
| Matrix size | N/A | N/A | 256 x 256 |
| **Functional** |  |  |  |
| Repetition time (TR) | 3480 ms | 2000 ms | 2130 ms |
| Echo time (TE) | 50 ms | 40 ms | 40 ms |
| Flip angle | 90 degrees | 90 degrees | 90 degrees |
| Voxel size | 3.8 x 3.8 x 5 mm | 4 x 4 x 5 mm | 4 x 4 x 5 mm |
| Matrix size | 64 x 64 | 64 x 64 | 64 x 64 |
| Number of slices | 27 | 28 | 32 |
| Field of view | N/A | 256 x 256 mm | 256 x 256 |
| Volumes [time] | 175 [8.05 min] | 239 [8 min] | 230 [8.17 min] |

### Neuroimaging Preprocessing

All three datasets were downloaded from their respective sources in BIDS format and were preprocessed using *fMRIPrep* version 1.2.6-1, an image processing pipeline based on *Nipype* 1.4.2 (Esteban et al., 2019; Gorgolewski et al., 2011; Gorgolewski et al., 2017). The details below were adapted from the fMRIPrep preprocessing details with extraneous details being omitted for clarity. Importantly, data from all datasets were preprocessed using the same pipeline.

The T1w image was corrected for intensity non-uniformity (INU) with N4BiasFieldCorrection (Tustison et al. 2010), distributed with ANTs 2.2.0 (Avants et al. 2008), and used as T1w-reference throughout the workflow. Skull-stripping was then performed on the T1w-reference using a *Nipype* implementation of the antsBrainExtraction.sh workflow (from ANTs), with OASIS30ANTs as the target template. Brain tissue segmentation of cerebrospinal fluid (CSF), white-matter (WM) and gray-matter (GM) was performed on the brain-extracted T1w using fast (FSL 5.0.9, Zhang, Brady, and Smith 2001). Volume-based spatial normalization to MNI152NLin2009cAsym standard space was performed through nonlinear registration with antsRegistration (ANTs 2.2.0), using brain-extracted versions of both T1w reference and the T1w template. To this end, the *ICBM 152 Nonlinear Asymmetrical template version 2009c* (Fonov et al. 2009) template was selected for spatial normalization.

For each of the BOLD runs contained per subject, the following preprocessing steps were performed. First, a reference volume and its skull-stripped version were generated using a custom methodology of fMRIPrep. Head-motion parameters with respect to the BOLD reference (transformation matrices, and six corresponding rotation and translation parameters) were estimated before any spatiotemporal filtering using mcflirt (FSL 5.0.9, Jenkinson et al. 2002). BOLD runs were slice-time corrected using 3dTshift from AFNI 20160207 (Cox 1996; Cox and Hyde 1997). Based on the estimated susceptibility distortion, a corrected EPI reference was calculated for a more accurate co-registration with the anatomical reference. The BOLD reference was then co-registered to the T1w reference using flirt (FSL 5.0.9, Jenkinson and Smith 2001) with the boundary-based registration (Greve and Fischl 2009) cost-function. Co-registration was configured with nine degrees of freedom to account for distortions remaining in the BOLD reference. The BOLD time-series (including slice-timing correction when applied) were resampled onto their original, native space by applying a single, composite transform to correct for head-motion and susceptibility distortions. These resampled BOLD time-series will be referred to as preprocessed BOLD in original space, or just preprocessed BOLD. The BOLD time-series were resampled into standard space, generating a preprocessed BOLD run in MNI152NLin2009cAsym space.

Automatic identification of motion artifacts using independent component analysis (ICA-AROMA, Pruim et al. 2015) was performed on the preprocessed BOLD on MNI space time-series after removal of non-steady state volumes and spatial smoothing with an isotropic, Gaussian kernel of 6mm FWHM (full-width half-maximum). AROMA motion components were subsequently included as regressors in our analyses (see below). Additional confounding time-series were calculated based on the preprocessed BOLD: framewise displacement (FD) and three regional signals (cerebral spinal fluid, white matter, and grey matter). FD was computed using the relative root mean square displacement between affines (Jenkinson et al. 2002). The three global signals are extracted within the CSF, the WM, and the whole-brain masks. Although, we note that FD was not used for “scrubbing” (Power et al., 2012; Power et al., 2014). All resamplings were performed with a single interpolation step by composing all the pertinent transformations (i.e., head-motion transform matrices, susceptibility distortion correction when available, and co-registrations to anatomical and output spaces). Gridded (volumetric) resamplings were performed using antsApplyTransforms (ANTs), configured with Lanczos interpolation to minimize the smoothing effects of other kernels (Lanczos 1964).

We removed subjects based on the Entropy Focus criterion (efc), Foreground to Background Energy Ratio (fber), Temporal Signal to Noise Ratio (tsnr), average framewise displacement (fd_mean), and Ghost to Signal Ratio in the Y direction (gsr_y) Image Quality Metrics (IQMs) from MRIQC (Esteban, et al., 2017). Outlier runs were defined as runs with efc, fd_mean, or gsr_y values exceeding 1.5 times the inter-quartile range above the 75th percentile, as well as those with fber and tsnr values lower than 1.5 times the lower bound minus the 25th percentile (i.e., a boxplot threshold). Using these parameters, 8 subjects were excluded for a total sample of 103 participants: 55 PD patients and 48 healthy controls.

### First-level Neuroimaging Analysis

Whole-brain analysis was performed across the three datasets using the FMRI Expert Analysis Tool (FEAT) from the FMRIB Software Library (FSL) to determine differences in functional connectivity of midbrain regions between healthy controls and PD patients. Seed regions, including the substantia nigra (SN) and ventral tegmental area (VTA), were defined using probabilistic atlases from previous work (Murty, et al., 2014). These atlases were developed by averaging 50 hand-drawn ROIs based on previous methods utilizing anatomical landmarks in the midbrain, which allowed for the separation of the SN and VTA (Ballard et al., 2011; Naidich et al., 2009). These regions were then resampled to fit the dimensions of each of the 3 datasets using AFNI’s 3dResample (Figure 3; Cox, 1996).

**Figure 3.**
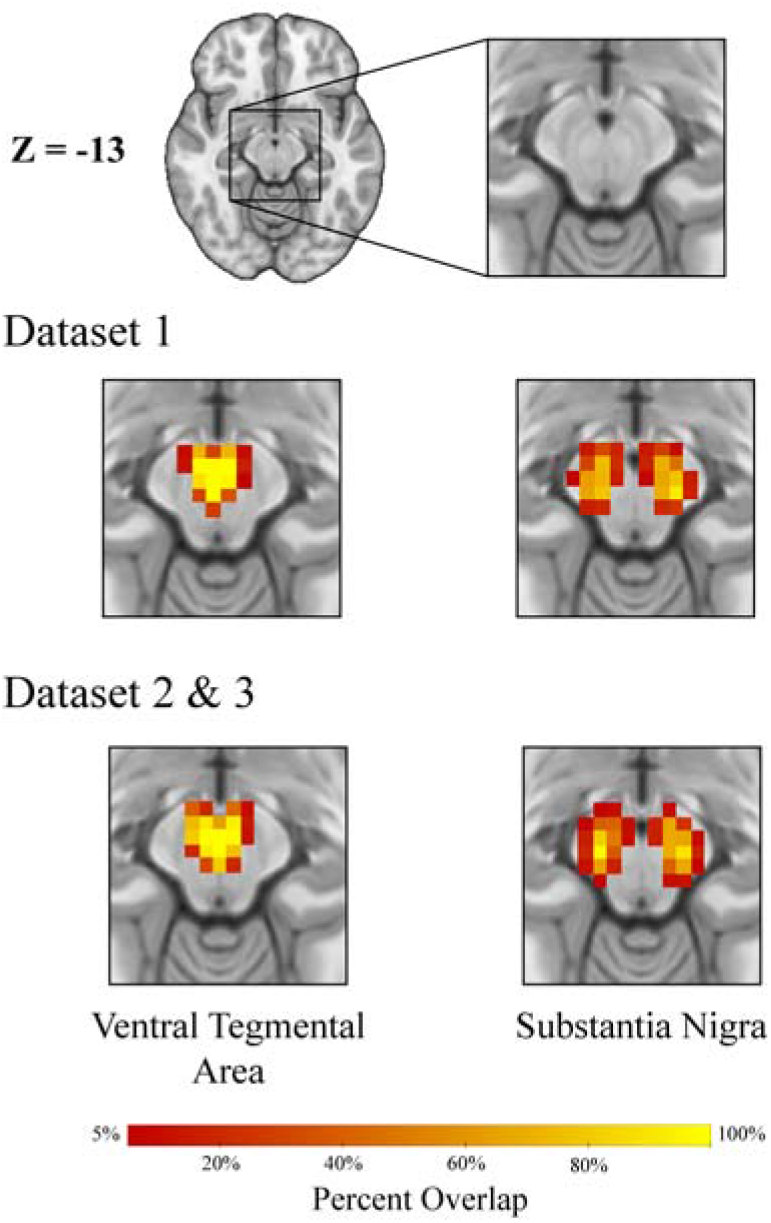
Seed ROIs for each dataset, based on probabilistic atlases (Murty et al., 2014). Dataset 1 has different voxel dimensions than datasets 2 and 3, therefore creating slightly different ROIs. Percent overlap refers to the probability that a voxel is overlapping with the anatomical demarcations of the ROI.

First, the eigenvariate of BOLD signal was extracted for each seed ROI within each subject using a weighted average based on each probabilistic atlas, which represents a single time series for each subject for each seed ROI. Then, FEAT was used to create a general linear model (GLM) with local correction for autocorrelation and regressors for the SN and VTA (Woolrich et al., 2001). Regression-based approaches are essential for estimating functional connectivity as Pearson correlations are much more likely to detect spurious relationships (Cole et al., 2019; Friston, 2011; Smith et al., 2016). This GLM created brain maps of functional connectivity within each subject for voxels predicted by the regressors of the following of contrasts of interest: SN > baseline, VTA > baseline, SN > VTA, and VTA > SN. The GLM also accounted for confound regressors of WM and CSF signal, non-steady-state volumes, cosine basis functions (to highpass filter with 128s cutoff), and ICA-AROMA motion. Importantly, functional connectivity cannot assess directionality. Indeed, a change in functional connectivity could reflect a change in signal in one region, a change in noise in another region, or a changes in connectivity with a third region (Friston, 2011; Smith et al., 2016).

### Higher-level Neuroimaging Analyses

For subjects of the NEUROCON dataset with two imaging runs, a second analysis was performed using a fixed effects model to create average BOLD maps across the runs for each subject. All brain maps were then used in a higher level random effects GLM analysis to compare functional connectivity of the entire sample across multiple regressors: group (control > PD, PD > control), ROI (SN > VTA, VTA > SN), and group by ROI interactions (Woolrich et al., 2004). This analysis utilized the Randomise function of FSL (Winkler et al., 2014) with Threshold-Free Cluster Enhancement (TFCE). All group-level analyses included covariates to control for potential confounds between groups and datasets. Importantly, the inclusion of dataset (dummy coded) and tsnr in our group-level analyses has been shown to control for variability in effect sizes in multisite studies (Friedman & Glover 2006). We further account for any differences in image quality across participants and datasets by including other image quality metrics in our model (i.e., efc, fber, fd_mean, and gsr_y). The data resolution is lower than is ideal for typical modern standards, however, based on our usage of confounding variables in our analyses, this should not affect our conclusions. Indeed, our models show that the site of data collection did not produce significant differences in the results of our whole-brain analysis. Finally, given that age (Varangis et al., 2019) and sex (Smith et al., 2014) have both been linked to differences in functional connectivity, these were also included as covariates in our analyses.

Any clusters of significant functional connectivity from the group by ROI interaction were further investigated by calculating an average value of functional connectivity across time points (t-stat) from both ROIs to that cluster in each subject. Then, two-tailed paired sample t-tests were run using R version 4.0.3 and RStudio (RStudio Team, 2020, version 1.3.1093; Allaire, 2012) to determine within-group and within-ROI effects. For the SN, we then used two-tailed paired sample t-tests to compare contralateral and ipsilateral connectivity to the clusters. Effects of laterality were not investigated in the VTA due to its medial location in the brain. For all statistical analyses, a significance threshold of p < 0.05 was used. For clusters in the cerebellum, a flat-map was created using the spatially unbiased atlas template of the cerebellum and brainstem (SUIT) toolbox (Diedrichsen & Zotow, 2015). The anatomical location of the clusters could then compared to the functional boundaries defined in recent work to infer its contributions to functioning (King et al., 2019). Any clusters were also input to LittleBrain to further interpret functional implications (Guell et al., 2019).

## Author Contributions

IMO performed data analysis, created figures, and drafted manuscript. HSP and IRO aided analysis and provided feedback for manuscript. VPM and DVS developed theoretical framework, initiated the project, and oversaw writing of the manuscript.

## Acknowledgments

We would like to thank the NEUROCON project, the Tao Wu group, and Tessa and colleagues for making their data publicly available. This work has been deposited on bioRxiv as a preprint. We note that DVS was a Research Fellow of the Public Policy Lab at Temple University during the preparation of this manuscript (2020 - 2021 academic year).

## References

Adcock, R. A., Thangavel, A., Whitfield-Gabrieli, S., Knutson, B., & Gabrieli, J. D. (2006). Reward-motivated learning: mesolimbic activation precedes memory formation. Neuron, 50(3), 507–517.

Allaire, J. (2012). RStudio: integrated development environment for R. Boston, MA, 770(394), 165–171.

Altmann, L. J., & Troche, M. S. (2011). High-level language production in Parkinson’s disease: a review. Parkinson’s disease, 2011.

Avants, B. B., Epstein, C. L., Grossman, M., & Gee, J. C. (2008). Symmetric diffeomorphic image registration with cross-correlation: Evaluating automated labeling of elderly and neurodegenerative brain. Medical Image Analysis, 12(1), 26–41. https://doi.org/10.1016/j.media.2007.06.004

Bäckman, L., Nyberg, L., Lindenberger, U., Li, S. C., & Farde, L. (2006). The correlative triad among aging, dopamine, and cognition: Current status and future prospects. In Neuroscience and Biobehavioral Reviews (Vol. 30, Issue 6, pp. 791–807). Pergamon. https://doi.org/10.1016/j.neubiorev.2006.06.005

Badea, L., Onu, M., Wu, T., Roceanu, A., & Bajenaru, O. (2017). Exploring the reproducibility of functional connectivity alterations in Parkinson’s disease. https://doi.org/10.1371/journal.pone.0188196

Ballard, I. C., Murty, V. P., Carter, R. M., MacInnes, J. J., Huettel, S. A., & Adcock, R. A. (2011). Dorsolateral prefrontal cortex drives mesolimbic dopaminergic regions to initiate motivated behavior. Journal of Neuroscience, 31(28), 10340–10346.

Berridge, K. C., Robinson, T. E., & Aldridge, J. W. (2009). Dissecting components of reward: “liking”, “wanting”, and learning. In Current Opinion in Pharmacology (Vol. 9, Issue 1, pp. 65– 73). Elsevier. https://doi.org/10.1016/j.coph.2008.12.014

Bromberg-Martin, E. S., Matsumoto, M., & Hikosaka, O. (2010). Dopamine in Motivational Control: Rewarding, Aversive, and Alerting. In Neuron (Vol. 68, Issue 5, pp. 815–834). Cell Press. https://doi.org/10.1016/j.neuron.2010.11.022

Buckner, R. L. (2013). The cerebellum and cognitive function: 25 years of insight from anatomy and neuroimaging. Neuron, 80(3), 807–815.

Caligiore, D., Arbib, M. A., Miall, R. C., & Baldassarre, G. (2019). The super-learning hypothesis: Integrating learning processes across cortex, cerebellum and basal ganglia. In Neuroscience and Biobehavioral Reviews (Vol. 100, pp. 19–34). Elsevier Ltd. https://doi.org/10.1016/j.neubiorev.2019.02.008

Carta, I., Chen, C. H., Schott, A. L., Dorizan, S., & Khodakhah, K. (2019). Cerebellar modulation of the reward circuitry and social behavior. Science, 363(6424). https://doi.org/10.1126/science.aav0581

Cole, M. W., Ito, T., Schultz, D., Mill, R., Chen, R., & Cocuzza, C. (2019). Task activations produce spurious but systematic inflation of task functional connectivity estimates. NeuroImage, 189, 1–18.

Cox, R. W. (1996). AFNI: Software for analysis and visualization of functional magnetic resonance neuroimages. Computers and Biomedical Research, 29(3), 162–173. https://doi.org/10.1006/cbmr.1996.0014

Cox, R. W., & Hyde, J. S. (1997). Software Tools for Analysis and Visualization of FMRI Data. NMR in Biomedicine, 10: 171–178

Dagher, A., & Robbins, T. W. (2009). Personality, Addiction, Dopamine: Insights from Parkinson’s Disease. In Neuron (Vol. 61, Issue 4, pp. 502–510). Cell Press. https://doi.org/10.1016/j.neuron.2009.01.031

D’Ambra, A. F., Jung, S. J., Ganesan S., Antzoulatos E. G., Fioravante D. (2021- submitted). Cerebellar Activation Bidirectionally Regulates Nucleus Accumbens Medial Shell and Core. bioRxiv.

Damier, P., Hirsch, E. C., Agid, Y., & Graybiel, A. M. (1999). The substantia nigra of the human brain II. Patterns of loss of dopamine-containing neurons in Parkinson’s disease. In Brain (Vol. 122).

Diedrichsen, J., & Zotow, E. (2015). Surface-Based Display of Volume-Averaged Cerebellar Imaging Data. https://doi.org/10.1371/journal.pone.0133402

Dreher, J. C., Meyer-Lindenberg, A., Kohn, P., & Berman, K. F. (2008). Age-related changes in midbrain dopaminergic regulation of the human reward system. Proceedings of the National Academy of Sciences, 105(39), 15106–15111.

Düzel, E., Bunzeck, N., Guitart-Masip, M., Wittmann, B., Schott, B. H., & Tobler, P. N. (2009). Functional imaging of the human dopaminergic midbrain. Trends in Neurosciences, 32(6), 321– 328. https://doi.org/10.1016/j.tins.2009.02.005

Esteban, O., Birman, D., Schaer, M., Koyejo, O. O., Poldrack, R. A., & Gorgolewski, K. J. (2017). MRIQC: Advancing the automatic prediction of image quality in MRI from unseen sites. https://doi.org/10.1371/journal.pone.0184661

Esteban, O., Markiewicz, C. J., Blair, R. W., Moodie, C. A., Isik, A. I., Erramuzpe, A., Kent, J. D., Goncalves, M., DuPre, E., Snyder, M., Oya, H., Ghosh, S. S., Wright, J., Durnez, J., Poldrack, R. A., & Gorgolewski, K. J. (2019). fMRIPrep: a robust preprocessing pipeline for functional MRI. Nature Methods, 16(1), 111–116. https://doi.org/10.1038/s41592-018-0235-4

Fearnley, J. M., & Lees, A. J. (1991). Ageing and parkinson’s disease: Substantia nigra regional selectivity. Brain, 114(5), 2283–2301. https://doi.org/10.1093/brain/114.5.2283

Fonov, V., Evans, A., McKinstry, R., Almli, C., & Collins, D. (2009). Unbiased nonlinear average age-appropriate brain templates from birth to adulthood. NeuroImage, 47, S102. https://doi.org/10.1016/s1053-8119(09)70884-5

Frazier, M., Hoffman L. J., Sullivan-Toole, H., Olino, T. M., Olson, I. R. (2021-submitted). A missing link in affect regulation: The cerebellum.

Friedman, L., Glover, G. H., & Fbirn Consortium. (2006). Reducing interscanner variability of activation in a multicenter fMRI study: controlling for signal-to-fluctuation-noise-ratio (SFNR) differences. Neuroimage, 33(2), 471–481.

Friston, K. J. (2011). Functional and effective connectivity: a review. Brain connectivity, 1(1), 13–36.

Goerendt, I. K., Lawrence, A. D., & Brooks, D. J. (2004). Reward Processing in Health and Parkinson’s Disease: Neural Organization and Reorganization. Cortex, 14, 73–80. https://doi.org/10.1093/cercor/bhg105

Gorgolewski, K., Burns, C. D., Madison, C., Clark, D., Halchenko, Y. O., Waskom, M. L., & Ghosh, S. S. (2011). Nipype: A flexible, lightweight and extensible neuroimaging data processing framework in Python. Frontiers in Neuroinformatics, 5. https://doi.org/10.3389/fninf.2011.00013

Gorgolewski, K. J., Esteban, O., Ellis, D. G., Notter, M. P., Ziegler, E., Johnson, H., Hamalainen, C., Yvernault, B., Burns, C., Manhães-Savio, A., Jarecka, D., Markiewicz, C. J., Salo, T., Clark, D., Waskom, M., Wong, J., Modat, M., Dewey, B. E., Clark, M. G., … Ghosh, S. (2017). Nipype: a flexible, lightweight and extensible neuroimaging data processing framework in Python. 0.13.1. https://doi.org/10.5281/ZENODO.581704

Gratton, C., Koller, J. M., Shannon, W., Greene, D. J., Maiti, B., Snyder, A. Z., Petersen, S.E., Perlmutter, J.S., & Campbell, M. C. (2019). Emergent functional network effects in Parkinson disease. Cerebral Cortex, 29(6), 2509–2523.

Greve, D. N., & Fischl, B. (2009). Accurate and robust brain image alignment using boundary-based registration. NeuroImage, 48(1), 63–72. https://doi.org/10.1016/j.neuroimage.2009.06.060

Guell, X., Goncalves, M., Kaczmarzyk, J. R., Gabrieli, J. D. E., Schmahmann, J. D., & Ghoshid, S. S. (2019). LittleBrain: A gradient-based tool for the topographical interpretation of cerebellar neuroimaging findings. https://doi.org/10.1371/journal.pone.0210028

Hacker, C. D., Perlmutter, J. S., Criswell, S. R., Ances, B. M., & Snyder, A. Z. (2012). Resting state functional connectivity of the striatum in Parkinson’s disease. Brain, 135(12), 3699–3711. https://doi.org/10.1093/brain/aws281

Hirsch, E., Graybielt, A. M., & Agid, Y. A. (1986). 28. National Research Council Nutrient Requirements of Beef Cattle Sixth revised edn Nutrient Requirements of Domestic Animals Number 4. In 16. Serengeti Ecological Monitoring Programme, Serengeti Wildlife Research Centre, PO Box (Vol. 17, Issue 2). Iowa State Univ. Press. Ames.

Id, C. T., Toschi, N., Id, S. O., Valenza, G., Lucetti, C., Barbieri, R., & Diciotti, S. (2019). Central modulation of parasympathetic outflow is impaired in de novo Parkinson’s disease patients. https://doi.org/10.1371/journal.pone.0210324

Ikai, Y., Takada, M., Shinonaga, Y., & Mizuno, N. (1992). Dopaminergic and non-dopaminergic neurons in the ventral tegmental area of the rat project, respectively, to the cerebellar cortex and deep cerebellar nuclei. Neuroscience, 51(3), 719–728.

Jenkinson, M. (2003). Fast, automated, N-dimensional phase-unwrapping algorithm. Magnetic Resonance in Medicine, 49(1), 193–197. https://doi.org/10.1002/mrm.10354

Jenkinson, M., Bannister, P., Brady, M., & Smith, S. (2002). Improved Optimization for the Robust and Accurate Linear Registration and Motion Correction of Brain Images. NeuroImage, 17(2), 825–841. https://doi.org/10.1006/nimg.2002.1132

Jenkinson, M., & Smith, S. (2001). A global optimisation method for robust affine registration of brain images. Medical Image Analysis, 5(2), 143–156. https://doi.org/10.1016/S1361-8415(01)00036-6

King, M., Hernandez-Castillo, C. R., Poldrack, R. A., Ivry, R. B., & Diedrichsen, J. (2019). Functional boundaries in the human cerebellum revealed by a multi-domain task battery. Nature Neuroscience, 22(8), 1371–1378. https://doi.org/10.1038/s41593-019-0436-x

Kwon, H. G., & Jang, S. H. (2014). Differences in neural connectivity between the substantia nigra and ventral tegmental area in the human brain. Frontiers in human neuroscience, 8, 41.

Lanczos, C. (1964). EVALUATION OF NOISY DATA*. In ANAL. Ser. B. https://epubs.siam.org/page/terms

Mallol, R., Barrós-Loscertales, A., López, M., Belloch, V., Parcet, M. A., & Ávila, C. (2007). Compensatory cortical mechanisms in Parkinson’s disease evidenced with fMRI during the performance of pre-learned sequential movements. Brain research, 1147, 265–271.

Milardi, D., Arrigo, A., Anastasi, G., Cacciola, A., Marino, S., Mormina, E., … & Quartarone, A. (2016). Extensive direct subcortical cerebellum-basal ganglia connections in human brain as revealed by constrained spherical deconvolution tractography. Frontiers in neuroanatomy, 10, 29.

Murty, V. P., Sambataro, F., Radulescu, E., Altamura, M., Iudicello, J., Zoltick, B., Weinberger, D.R., Goldberg, T.E. & Mattay, V. S. (2011). Selective updating of working memory content modulates meso-cortico-striatal activity. Neuroimage, 57(3), 1264–1272.

Murty, V. P., Shermohammed, M., Smith, D. V., Carter, R. M. K., Huettel, S. A., & Adcock, R. A. (2014). Resting state networks distinguish human ventral tegmental area from substantia nigra. NeuroImage, 100, 580–589. https://doi.org/10.1016/j.neuroimage.2014.06.047

Naidich, T. P., Duvernoy, H. M., Delman, B. N., Sorensen, A. G., Kollias, S. S., & Haacke, E. M. (2009). Duvernoy’s atlas of the human brain stem and cerebellum: high-field MRI, surface anatomy, internal structure, vascularization and 3 D sectional anatomy. Springer Science & Business Media.

Palmer, W. C., Cholerton, B. A., Zabetian, C. P., Montine, T. J., Grabowski, T. J., & Rane, S. (2021). Resting-State Cerebello-Cortical Dysfunction in Parkinson’s Disease. Frontiers in neurology, 11, 1855.

Power, J. D., Barnes, K. A., Snyder, A. Z., Schlaggar, B. L., & Petersen, S. E. (2012). Spurious but systematic correlations in functional connectivity MRI networks arise from subject motion. NeuroImage, 59(3), 2142–2154. https://doi.org/10.1016/j.neuroimage.2011.10.018

Power, J. D., Mitra, A., Laumann, T. O., Snyder, A. Z., Schlaggar, B. L., & Petersen, S. E. (2014). Methods to detect, characterize, and remove motion artifact in resting state fMRI. NeuroImage, 84, 320–341. https://doi.org/10.1016/j.neuroimage.2013.08.048

Pruim, R. H. R., Mennes, M., van Rooij, D., Llera, A., Buitelaar, J. K., & Beckmann, C. F. (2015). ICA-AROMA: A robust ICA-based strategy for removing motion artifacts from fMRI data. NeuroImage, 112, 267–277. https://doi.org/10.1016/j.neuroimage.2015.02.064

Reijnders, J. S. A. M., Ehrt, U., Weber, W. E. J., Aarsland, D., & Leentjens, A. F. G. (2008). A systematic review of prevalence studies of depression in Parkinson’s disease. In Movement Disorders (Vol. 23, Issue 2, pp. 183–189). https://doi.org/10.1002/mds.21803

Salamone, J. D., Correa, M., Farrar, A., & Mingote, S. M. (2007). Effort-related functions of nucleus accumbens dopamine and associated forebrain circuits. In Psychopharmacology (Vol. 191, Issue 3, pp. 461–482). Springer. https://doi.org/10.1007/s00213-006-0668-9

Sharman, M., Valabregue, R., Perlbarg, V., Marrakchi-Kacem, L., Vidailhet, M., Benali, H., Brice, A., & Lehéricy, S. (2013). Parkinson’s disease patients show reduced cortical-subcortical sensorimotor connectivity. Movement Disorders, 28(4), 447–454. https://doi.org/10.1002/mds.25255

Shohamy, D., & Wagner, A. D. (2008). Integrating memories in the human brain: hippocampal-midbrain encoding of overlapping events. Neuron, 60(2), 378–389.

Smith, D. V., Gseir, M., Speer, M. E., & Delgado, M. R. (2016). Toward a cumulative science of functional integration: A metalJanalysis of psychophysiological interactions. Human brain mapping, 37(8), 2904–2917

Smith, D. V., Utevsky, A. V., Bland, A. R., Clement, N., Clithero, J. A., Harsch, A. E., McKell Carter, R., & Huettel, S. A. (2014). Characterizing individual differences in functional connectivity using dual-regression and seed-based approaches. NeuroImage, 95, 1–12. https://doi.org/10.1016/j.neuroimage.2014.03.042

Stoodley, C. J., Valera, E. M., & Schmahmann, J. D. (2012). Functional topography of the cerebellum for motor and cognitive tasks: an fMRI study. Neuroimage, 59(2), 1560–1570.

Tessa, C., Toschi, N., Orsolini, S., Valenza, G., Lucetti, C., Barbieri, R., & Diciotti, S. (2019). Central modulation of parasympathetic outflow is impaired in de novo Parkinson’s disease patients. PloS one, 14(1), e0210324.

Tepfer, L. J., Alloy, L. B., & Smith, D. V. (2021). Family history of depression is associated with alterations in task-dependent connectivity between the cerebellum and ventromedial prefrontal cortex. Depression and Anxiety, 38(5). https://doi.org/10.1002/da.23143

Tomasi, D., & Volkow, N. D. (2014). Functional connectivity of substantia nigra and ventral tegmental area: maturation during adolescence and effects of ADHD. Cerebral cortex, 24(4), 935–944.

Tustison, N. J., Avants, B. B., Cook, P. A., Zheng, Y., Egan, A., Yushkevich, P. A., & Gee, J. C. (2010). N4ITK: Improved N3 bias correction. IEEE Transactions on Medical Imaging, 29(6), 1310–1320. https://doi.org/10.1109/TMI.2010.2046908

Varangis, E., Habeck, C. G., Razlighi, Q. R., & Stern, Y. (2019). The effect of aging on resting state connectivity of predefined networks in the brain. Frontiers in aging neuroscience, 11, 234.

Wei, L., Hu, X., Yuan, Y., Liu, W., & Chen, H. (2018). Abnormal ventral tegmental area-anterior cingulate cortex connectivity in Parkinson’s disease with depression. Behavioural brain research, 347, 132–139.

Winkler A. M., Ridgway G. R., Webster M. A., Smith S. M., Nichols T. E. (2014). Permutation inference for the general linear model. NeuroImage, 2014;92:381–397

Wise, R. A. (2004). Dopamine, learning and motivation. In Nature Reviews Neuroscience (Vol. 5, Issue 6, pp. 483–494). Nature Publishing Group. https://doi.org/10.1038/nrn1406

Woolrich, M. W., Behrens, T. E. J., Beckmann, C. F., Jenkinson, M., & Smith, S. M. (2004). Multilevel linear modelling for FMRI group analysis using Bayesian inference. NeuroImage, 21(4), 1732–1747. https://doi.org/10.1016/j.neuroimage.2003.12.023

Woolrich, M. W., Ripley, B. D., Brady, M., & Smith, S. M. (2001). Temporal autocorrelation in univariate linear modeling of FMRI data. Neuroimage, 14(6), 1370–1386.

Wu, T., & Hallett, M. (2013). The cerebellum in Parkinson’s disease. In Brain (Vol. 136, Issue 3, pp. 696–709). Oxford University Press. https://doi.org/10.1093/brain/aws360

Wu, T., Wang, J., Wang, C., Hallett, M., Zang, Y., Wu, X., & Chan, P. (2012). Basal ganglia circuits changes in Parkinson’s disease patients. Neuroscience letters, 524(1), 55–59.

Zhang, H. Y., Tang, H., Chen, W. X., Ji, G. J., Ye, J., Wang, N., Wu, J.T., & Guan, B. (2015). Mapping the functional connectivity of the substantia nigra, red nucleus and dentate nucleus: a network analysis hypothesis associated with the extrapyramidal system. Neuroscience letters, 606, 36–41.

Zhang, S., Hu, S., Chao, H. H., & Li, C. S. R. (2016). Resting-state functional connectivity of the locus coeruleus in humans: in comparison with the ventral tegmental area/substantia nigra pars compacta and the effects of age. Cerebral Cortex, 26(8), 3413–3427.

Zhang, Y., Brady, M., & Smith, S. (2001). Segmentation of brain MR images through a hidden Markov random field model and the expectation-maximization algorithm. IEEE Transactions on Medical Imaging, 20(1), 45–57. https://doi.org/10.1109/42.906424

