## Supplementary Information for "Distinct Alterations in Cerebellar Connectivity with Substantia Nigra and Ventral Tegmental Area in Parkinson’s Disease"

***Affiliations***
[a-e]: Department of Psychology, Weiss Hall, Temple University, 1701 N. 13^th^ St, Philadelphia, PA 19112, USA

***Correspondence***
David V. Smith or Vishnu P. Murty
Department of Psychology, Temple University.

**Supplementary Methods:**

*Cerebellar laterality analysis*

To assess laterality of midbrain connectivity with the cerebellum, it was first necessary to left-right flip our cerebellar ROI using FSL (Jenkinson et al., 2012). For both right and left cerebellar ROIs, an average value of functional connectivity across time points (t-stat) from both seed regions was calculated for each subject. We then used RStudio to fit a mixed effects linear regression model with terms for seed roi, group, region laterality, and their interaction terms to predict connectivity in the cerebellum (RStudio Team, 2020, version 1.3.1093). The model also account for the random effects of subjects.

*Effects of age on connectivity*

As an exploratory analysis, we tested correlations between age and midbrain connectivity in our sample. We used the *cor.test* function in RStudio to dertermine if increases in age predicted SN or VTA connectivity to regions of interest including the caudate, putamen, nucleus accumbens, and cerebellum (RStudio Team, 2020, version 1.3.1093).

*Midbrain interactions with striatum*

Following our original analysis, we were interested in any interaction of the VTA and SN with striatal regions. To investigate any differences in connectivity, we calculated average connectivity of VTA and SN with striatal regions including the caudate, putamen, and nucleus accumbens (NAcc). We used RStudio to run a 2 * 2 ANOVA to test for effects of seed ROI, group, and their interaction in each striatal ROI. (RStudio Team, 2020, version 1.3.1093)

**Supplementary Results:**

In our analysis of the laterality of cerebellar-midbrain connectivity, we found a three-way interaction such that the seed roi * group effect was significantly stronger in the right cerebellum than the left (F(1, 404) = 12.4, p < 0.001). In our analysis of the effects of age on connectivity, we found a strending negative correlation between age and connectivity between the SN and caudate (r = -0.18, p = 0.066). Finally, we found no effects of seed ROI (caudate: F(1, 202) = 0.47, p = 0.493; putamen: F(1, 202) = 1.24, p = 0.266, NAcc: F(1, 202) = 0.78 p = 0.381), group (caudate: F(1, 202) = 0.04 p = 0.842; putamen: F(1, 202) = 0.66 p = 0.418, NAcc: F(1, 202) = 0 p = 0.996), or their interaction (caudate: F(1, 202) = 1.20, p = 0.275; putamen: F(1, 202) = 1.96, p = 0.163, NAcc: F(1, 202) = 2.08, p = 0.151) on connectivity of the midbrain with any striatal regions.

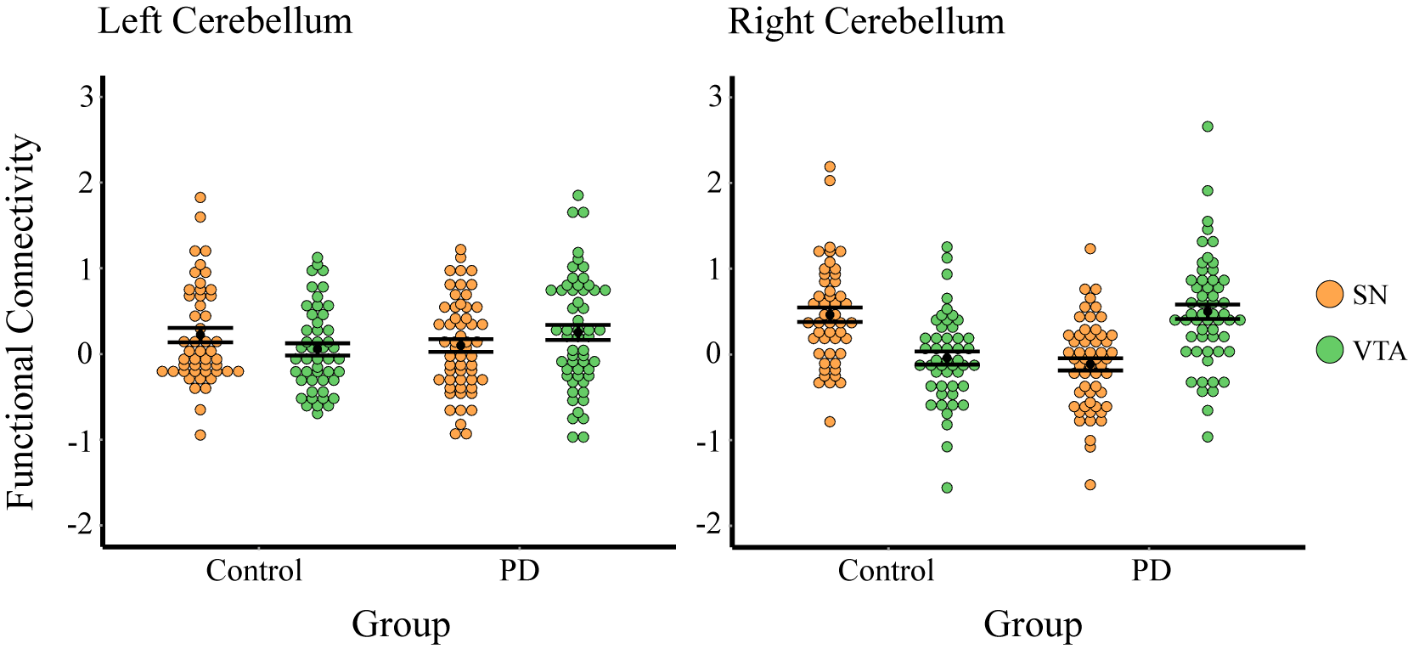

Supplementary figure 1. Dotplot shows that the right cerebellum demonstrates a stronger effect of the seed roi * group interaction in midbrain connectivity.

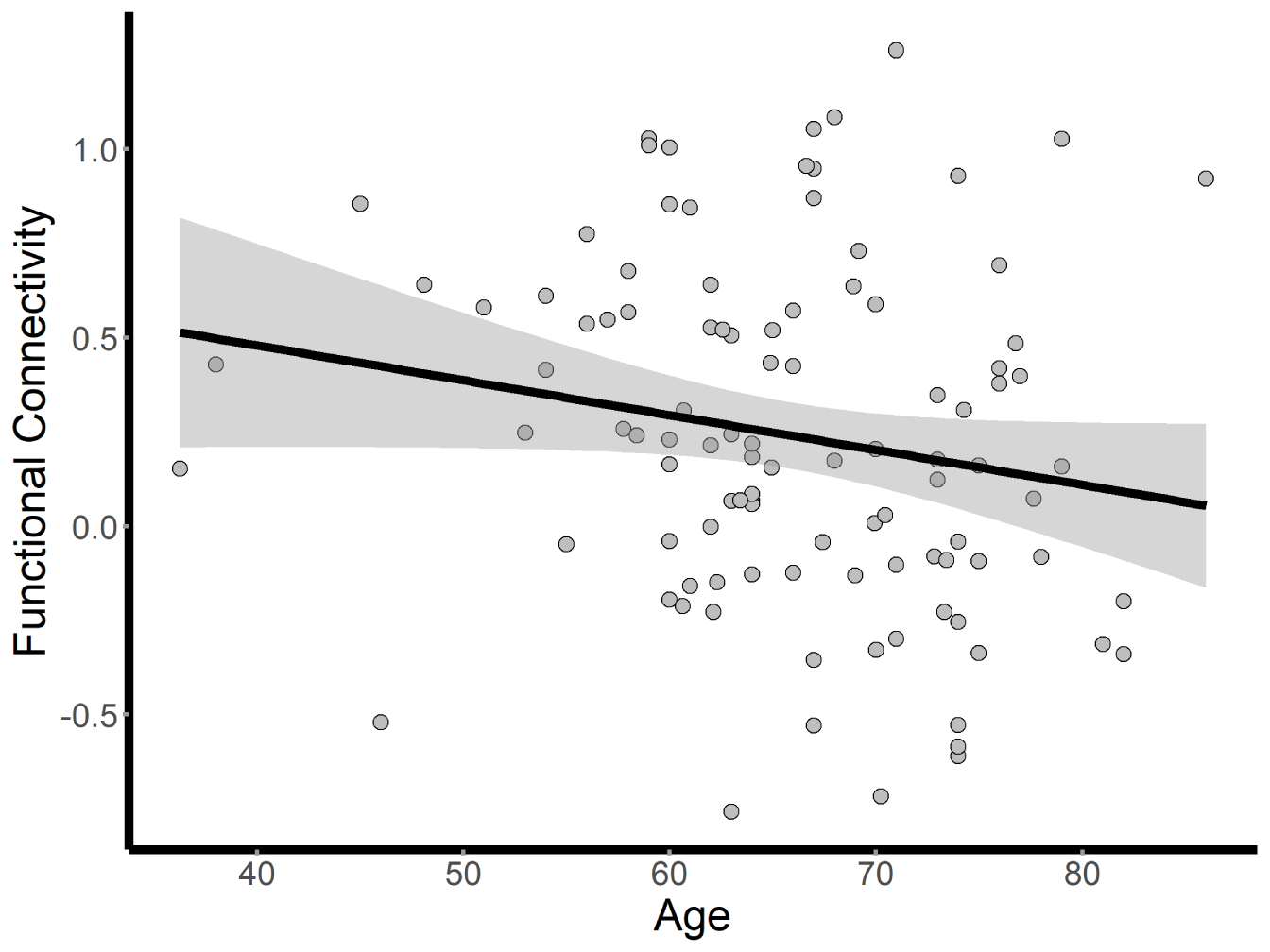

Supplementary figure 2. Scatter plot shows a trending negative correlation between age of participants and connectivity between SN and caudate.

Works Cited:

Allaire, J. (2012). RStudio: integrated development environment for R. *Boston, MA*, *770*(394), 165-171.

Jenkinson, M., Beckmann, C. F., Behrens, T. E., Woolrich, M. W., & Smith, S. M. (2012). FSL. *Neuroimage*, *62*(2), 782-790.
